# Scalable *in-vitro* immunostimulation of human blood for in-depth profiling of acute immune response effects

**DOI:** 10.1101/2023.06.08.544193

**Authors:** Stefan Markus Reitzner, Petter Brodin, Jaromír Mikeš

## Abstract

In-vitro immune stimulation of whole blood has great analytic potential for exploring the immune system function. However, logistical and biological constraints might limit the extent to which such investigations can be performed. Flexible and often even low-tech mobile applicability can increase the diversity of conditions that can be investigated. To this end, we developed and optimized a medium that enables prolonged *in-vitro* stimulation and enables highly functional immune responses within 24 hours of incubation. In addition, we also developed and optimized a low-tech water-based mobile incubator to enable logistical flexibility. Finally, we describe the implementation of both systems using a practical example of sample collection at a sampling site with no access to laboratory equipment outside of a research lab.

## Background

In-vitro immune stimulation of whole blood using pathogens has emerged as a useful tool to study immunocompetence and molecular mechanisms of immune cell activation in a variety of human conditions and settings^1–3^. Briefly, a freshly sampled blood supplemented with a nutrient medium is being exposed to stimulants such as Influenza A virus (IAV), bacterial lipopolysaccharide (LPS), Bacillus Calmette-Guérin (BCG) or similar, constituting an immune challenge. Technical details, such as the incubation period, can differ depending on the requirements of the experimental design or the investigated effect. Even though this is a straightforward approach, the need for specialized lab equipment significantly limits the range of options and requires proximity to lab facilities. This, however, excludes more specialized or extreme conditions, which for the same reason, might be more valuable to explore. To this end, we developed several methods and techniques enabling a mobile implementation of immuno-monitoring methods, including an easy-to-use low-tech mobile incubator and a suitable incubation medium.

### Development of a stimulation medium

First, to address the survival of whole blood-born immune cells in a stimulation reaction, we had to develop a medium that can provide an appropriate environment and sufficient supply of nutrients to endure prolonged *in-vitro* stimulation periods and immune challenges with high viability and RNA quality. To do so, we conducted several pilot experiments. We tested cell survival and RNA quality under different conditions to identify key components having a positive or negative impact on cell survival within a prolonged incubation period. Excluding some of the common additives, such as bovine fetal serum, we created two candidate media, A and B, with a proprietary cocktail of additives which, based on gene set enrichment analysis (GSEA), improved the functional immune cell response based on their activated pathways compared to incubation without it (Figure 1A). In further experiments using LPS as an immune stimulant, we identified our prepared medium A with the cocktail to be superior to medium B as measured by GSEA functional immune response (Figure 1B,C). Based on these findings, we developed the IAMTHESTIM (Cytodelics AB, Stockholm, Sweden) stimulation medium, which was subsequently tested with the three different stimulants, IAV, LPS, and BCG for up to 7 days (Fig. 2). We found high viability within 48 hours of incubation, good quality (RIN between 6.3 and 6.6), and quantity of RNA (between 4279 and 3288pg/μl) using the PAXgene RNA extraction system. Following seven days of incubation, RNA quantity decreased by a factor of 6 to 7, but the quality dropped only marginally reflecting the viability between 30 and 40%. Yet, this RNA was still of satisfying quality and suitable for downstream analyses, demonstrating the potential of this stimulation system.

**Figure 1:**
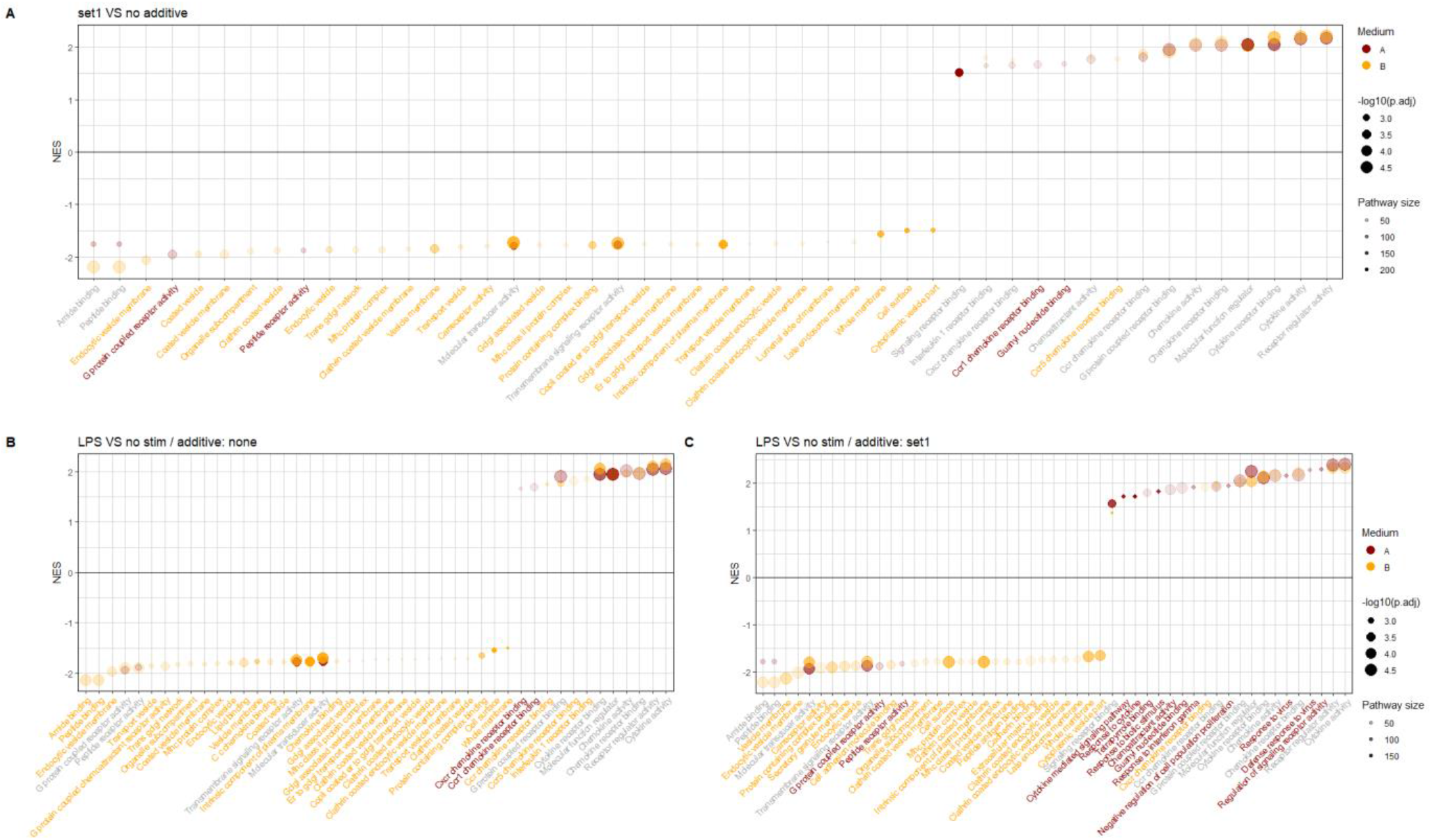
Functional analysis of different media and additives in response to no stimulation and LPS stimulation following 21 hours of immune stimulation. (A) Comparing set1 additives to no additives with two different candidate media without immune stimulation. (B) Comparing LPS stimulation with no stimulation in a medium without additive in two different candidate media. (C) Comparing LPS stimulation with no stimulation in a medium with set1 additives in two different candidate media.

**Figure 2:**
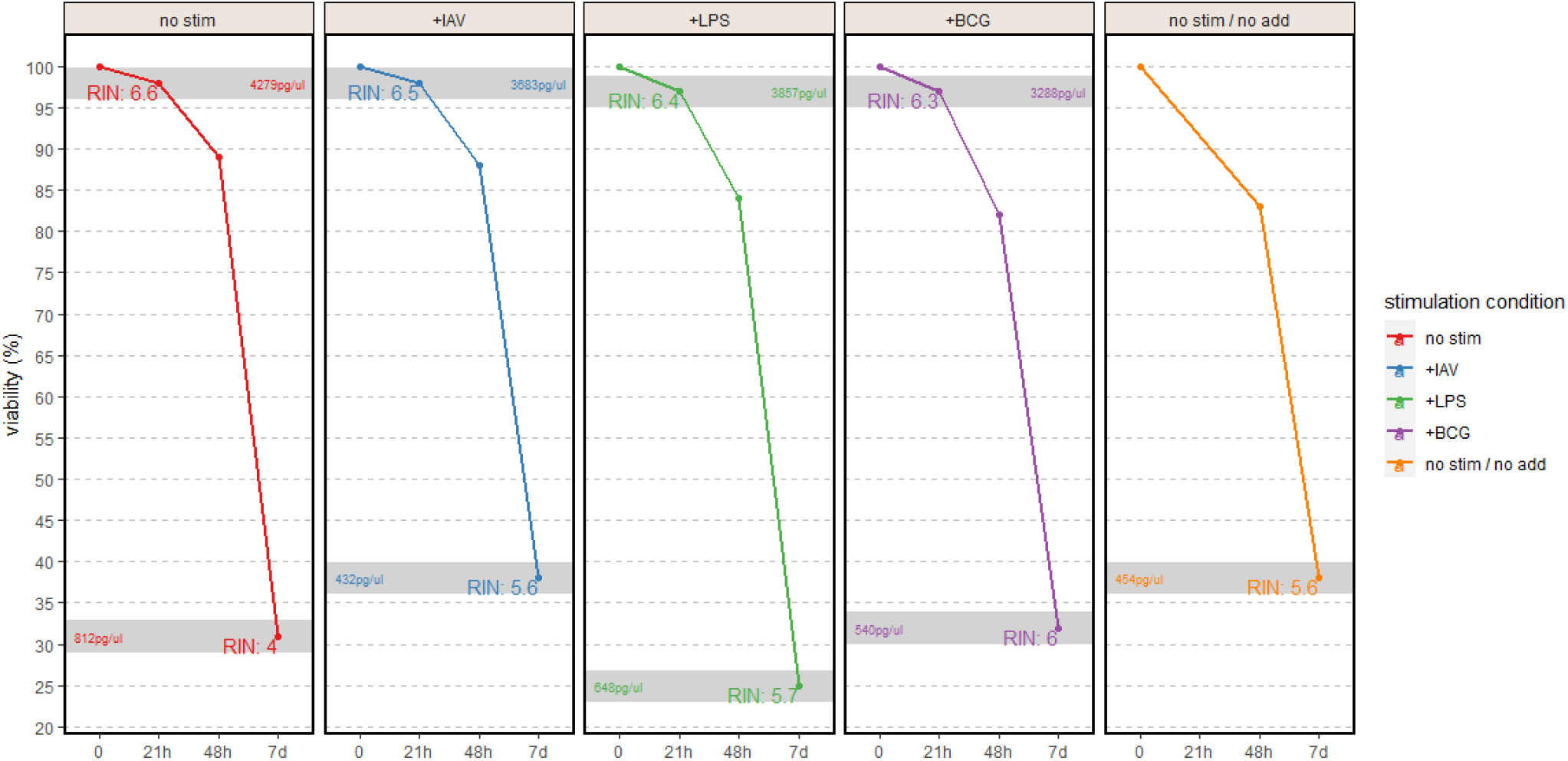
Cell viability and RNA quality Samples with different stimulants (IAV, LPS or BCG) or with a cell viability increasing additive were incubated for 21 and 48 hours as well as 7 days, their viability measured using the Cellaca system and their RNA quality quantified with a Qubit fluorometer.

### Development of a mobile incubation protocol and system

Utilization of the developed stimulation system in a flexible, non-stationary environment, requires relatively constant incubation temperature in a physiological range from 34.5°C to 37.5°C for a relatively long period of time. Therefore, we tested various approaches to provide such conditions. First, we were considering a series of dry baths using battery power in a system involving a car battery or alternator during the transport period. However, commercially available options did not match the available maximal power output capacity in standard personal vehicles considering our requirements for the number of samples to be collected. Next, we were considering isothermal delta-phase gel pads that were designed to maintain 37°C. Unfortunately, a test of this setup in a Styrofoam box revealed low thermal stability, with the usable time window being only 2 hours and 10 minutes long (Fig. 3A). Instead, we changed our approach to large volumes of water. For this, we used five double-zip lock bags, each filled with water to half their nominal volume, 16L in total. A large Styrofoam box was used with a wall thickness of 50 mm and a total internal volume of about 30 dm^3^ (31 × 31 × 31 cm) for isolation. The water volume in the zip lock bags was heated overnight to 39°C a day before the experiment. This resulted in an 8 hours and 25 minutes long time window with the temperature within the parameters of the assay (Fig. 3B). Additionally, this setup provides us with a 4 hours 55 minutes buffer to conduct our experiment on site and collect the blood samples before the useable time window starts. Altogether, these parameters satisfied and significantly exceeded the minimal requirements set for the mobile incubation solution in the context of our project.

**Figure 3:**
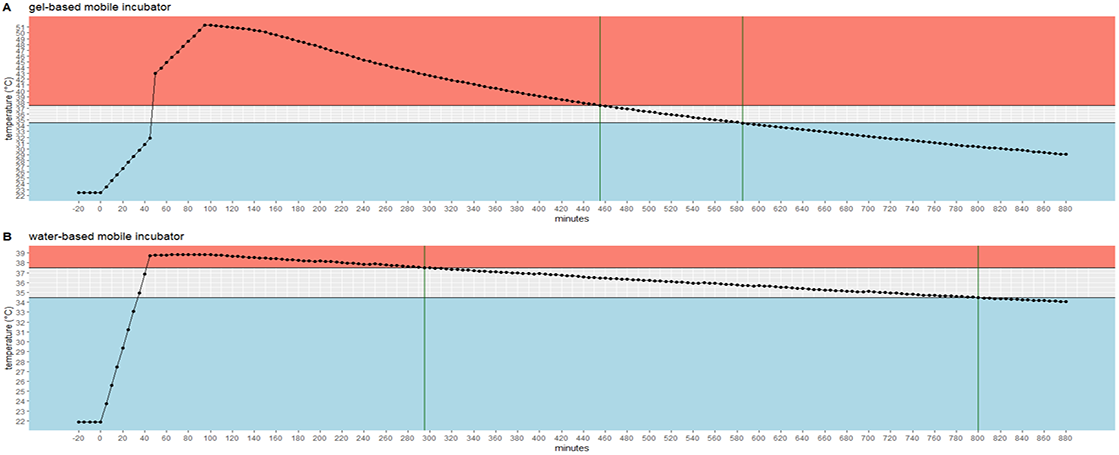
Temperature stability of tested mobile incubation systems Water-and gel-based mobile Styrofoam incubator performance to reach a target temperature of 34.5-37.5°C for a maximized duration to enable a mobile, power-less incubation of stimulation samples.

### Specific implementation of medium and incubation system in an on-site acute blood stimulation experiment

Following the development and quality control of the incubation medium chemistry and the low-tech mobile incubation solution, we implemented these elements into an on-site acute exercise stimulation experiment for our ongoing ‘iSOK’ study. In particular, we wanted to investigate the response of the circulating immune system to three different immune challenges (5ng/ml LPS, 50 HAU IAV, and BCG (2-8•10^5^ BAC per stimulation) and medium only control at five time points around a roughly 1.5 hours long acute bout of exercise in twelve athletes. This resulted in 20 stimulation vials per subject and 240 stimulation vials for the whole experiment. Three 80-vial rack blocks were used to fit them into the mobile incubator design in between the water bags (Fig. 4).

**Figure 4:**
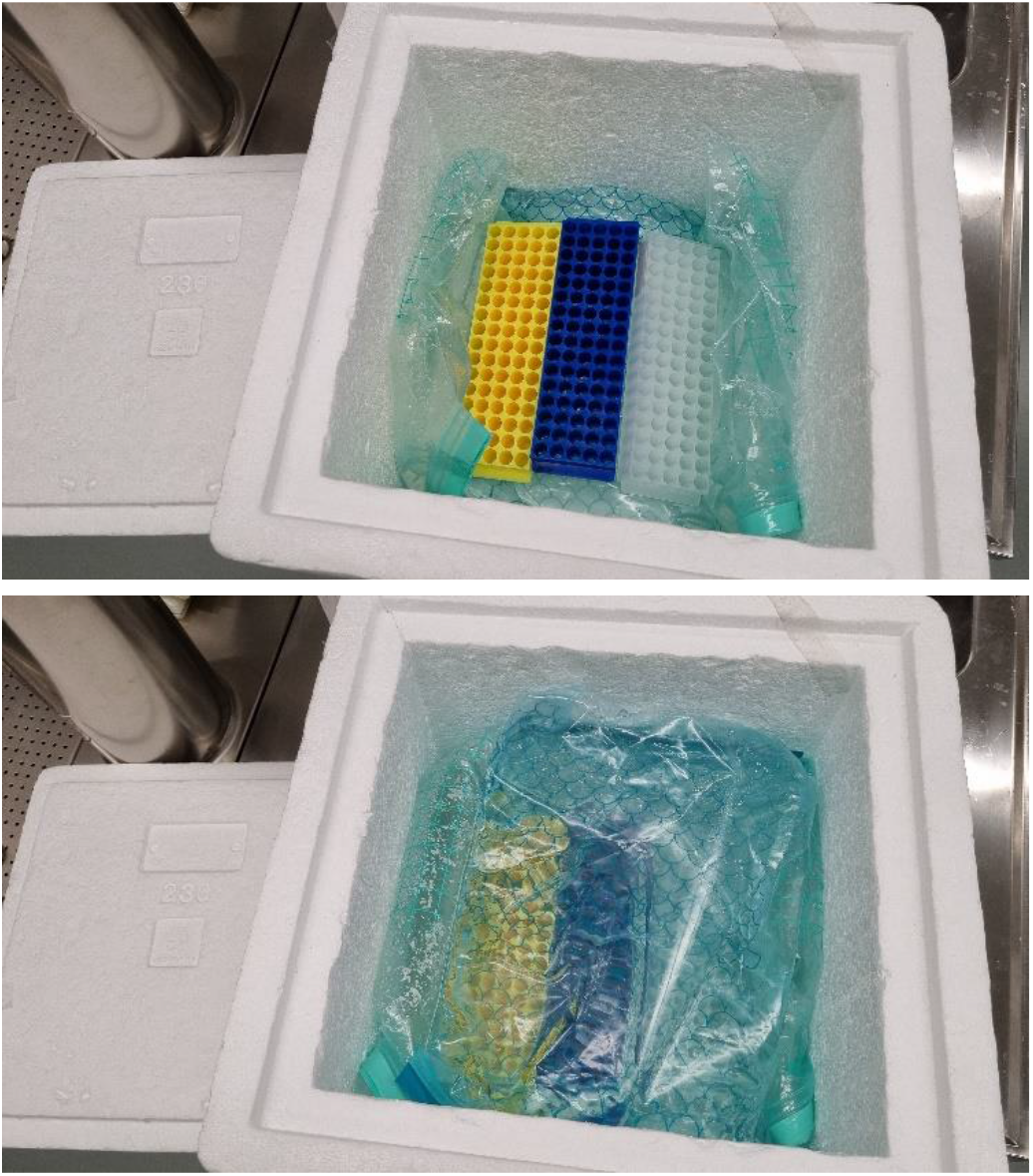
Mobile incubator low-tech solution using water as a temperature stabilizer for keeping an incubation temperature between 34.5 and 37.5°C for 240 stimulations in 2ml vials.

One day before intervention day, the zip-lock bags with water were placed in an incubator set to 39°C overnight. On the morning of the intervention day, they were packed into the Styrofoam box with the rack blocks and a simple aquarium temperature sensor that displays the current temperature as well as minimum and maximum temperature to ensure the quality of the temperature environment. The prepared incubation vials were transported on wet ice in a separate box. On-site, venous blood was collected from each of the 12 subjects before acute exercise, 1ml each immediately stimulated in the prepared stimulation vials, and the incubation timer started. Additionally, without any stimulation, 500μl of whole blood was stabilized using the Cytodelics whole blood stabilization system (Cytodelics AB, Stockholm, Sweden), and 1ml whole blood was treated with 2ml PAXgene solution (Becton Dickinson, Franklin Lakes, NJ, USA). The remaining collected whole blood was stored on ice. After the research subjects performed their acute exercise, blood sampling and stimulation were repeated immediately after and 30 minutes, 1 hour and 2 hours after the end of acute exercise. After the collection was completed and 240 stimulations in total were initiated using the mobile incubator, all samples were transferred to the laboratory. The stimulation continued in an incubator set to 37°C/5% CO_2_/95% humidity for the remaining time, in total 21-24 hours. All tube screw-caps were slightly opened to allow CO_2_ exchange with the buffer system in the stimulation medium. Remaining non-stimulated whole blood was centrifuged at 3000g for 10 minutes, plasma collected and frozen at -80°C together with the PAXgene-preserved whole blood collected during the day.

On the next day, at the same time as the stimulation timer was started, starting with the samples collected before the acute exercise was performed, samples were properly resuspended, and 200μl of the blood and stimulation buffer mixture was transferred into prepared cryovials with 200μl Cytodelics Stabilizer, incubated at room temperature for 10 minutes and frozen at -80°C. The remaining volume was centrifuged (1200g, 5 minutes, RT), and the plasma was collected and frozen at -80°C. The cell layer was mixed with 2ml of PAXgene solution, transferred into 4ml cryovials, incubated (2 hours, RT), and finally frozen at -80°C. This was repeated in chronological order for the other stimulations sampling time points.

The final product of this collection day was 75 samples for each subject; 60 stimulated and 15 non-stimulated; 25 immune cell samples, 25 plasma samples, and 25 PAXgene lysed and stabilized RNA samples stored at -80°C. In total, 900 samples were produced as a result of one sample field collection trip.

The described sampling procedure was repeated until study completion. Immune cells cryogenically preserved in the Cytodelics Stabilizer and extracted using the Cytodelics whole blood processing system were used for in-depth mass cytometric cell population analysis in response to acute exercise, enabling us to understand the dynamics of cell expansion or recruitment. Gene expression changes in immune cells in response to acute exercise were analyzed from bulk-sequenced RNA samples extracted from PAXgene-preserved samples. Cytokines released from immune cells were detected by plasma proteomic analysis from the collected plasma samples. Taken together, this extensive but technically simple approach enabled a comprehensive, system-wide analysis of the activities of the immune system in response to acute exercise.

